# AI-Guided Dual Strategy for Peptide Inhibitor Design Targeting Structural Polymorphs of α-Synuclein Fibrils

**DOI:** 10.1101/2025.10.10.681708

**Authors:** Jinfang Duan, Haoyu Zhang, Chuanqi Sun

## Abstract

One of the most important events in the pathogenesis of Parkinson’s disease and related disorders is the formation of abnormal fibrils via the aggregation of α-synuclein (α-syn) with β-sheet-rich organization. The use of Cryo-EM has uncovered different polymorphs of the fibrils, each having unique structural interfaces, which has made the design of inhibitors even more challenging. Here, a structure-guided framework incorporating AI-assisted peptide generation was set up with the objective of targeting the conserved β-sheet motifs that are present in various forms of α-syn fibrils. The ProteinMPNN, then, AlphaFold-Multimer, and PepMLM were employed to create short peptides that would interfere with the growth of the fibrils. The two selected candidates, T1 and S1, showed a significant inhibition of α-syn fibrillation, as measured by a decrease in the ThT fluorescence and the generation of either amorphous or fragmented aggregates. The inhibitory potency of the peptides was in line with the predicted interface energies. This research work illustrates that the integration of cryo-EM structural knowledge with the computational design method leads to the quick discovery of the wide-spectrum peptide inhibitors, which is a good strategy for the precision treatment of neurodegenerative diseases.

## 1. Introduction

Parkinson’s disease (PD) stands as the second largest among the neurodegenerative disorders that affect people all over the world. It is mainly characterized by the gradual deterioration of motor function, the death of dopaminergic neurons in the brain region called substantia nigra, and the formation of abnormal intracellular aggregations known as Lewy bodies and Lewy neurites[1-3]. The major protein that forms these inclusions is α-synuclein (α-syn), a 140-amino-acid presynaptic protein that switches from a dissolved, naturally disordered monomer to insoluble β-sheet-rich amyloid fibrils[4-7]. Such pathological conversion is a characteristic of PD and related synucleinopathies like MSA and DLB[6,8,9].

Cryo-electron microscopy (cryo-EM) has recently made significant progress in revealing that not all α-syn fibrils are of the same kind but rather exist as a whole range of structural polymorphs, each characterized by specific protofilament arrangements, interfacial salt-bridge networks, and β-sheet stacking patterns[10,11]. The different types of this polymorphic fibril have been recognized in biopsies from patients, amplified by seeding, and in recombinant assemblies, showing both disease-specific and common structural features[12-14]. Moreover, the different types of fibrils have been found to be biologically active in different ways, hence causing differences in seeding potency, cytotoxicity, and ultimately, disease phenotype [15,16]. One could rightly say that this heterogeneity in structure is the main reason behind the different clinical presentations seen in synucleinopathies. The various forms of α-syn fibrils can be considered as a major hurdle in the way of the development of new treatments [17]. A lot of small molecules and antibodies deal with the problem by specifically recognizing the different conformations or the corresponding regions of the molecule and hence being unable to stop the action of the other strains of fibrils[18,19]. In addition, it is true that the majority of the currently available inhibitors work only at the very beginning stages of oligomerization and therefore the whole process of fibril elongation remains more or less unhampered[20]. Researchers have found out that peptide-based inhibitors can bring about an enormous change and have become the most sought-after in the drug industry owing to their ability to provide very high structural specificity, chemical tunability, and mimicking of natural protein interfaces[21,22]. There is a possibility for short peptides to be designed in such a way as to bind with the aggregation-prone β-sheet motifs, cap the fibril ends, or even destabilize the interactions between the protofilaments. But the enormous variety of possible sequences of peptides that can be combined together and the structural diversity of fibrils make it very hard to find the right peptide sequences which are both effective and have broad inhibitory activity.

Over the past few years, artificial intelligence (AI) has completely changed the field of protein and peptide design [23-25]. Among them, the strongest algorithms for structure prediction, like AlphaFold, as well as generative sequence models that have been trained with very large peptide datasets, nowadays allow to rapidly scan huge sequence spaces with accuracy down to the atomic level[22,26]. The same techniques can be employed to design the inhibitors that can bind to the specified aggregation motifs, and at the same time, be the most stable and soluble ones through physicochemical property optimization[25]. The ability to work with high-resolution cryo-EM templates along with AI revolutionized design and made it a powerful and efficient route for the rational development of peptide inhibitors targeting amyloid fibril polymorphs[27-29].

In this study, a dual AI-guided design framework was created for the generation of peptide inhibitors that target α-syn fibrils. Initially, the research group mapped out the conserved β-sheet motifs over various cryo-EM-resolved polymorphs and flagged them as the universal inhibitory targets[1,3,5,8,11,30-43]. Computational methods were then applied in two ways: firstly, Structure-based design; in this case, ProteinMPNN made the selection of peptides fitting into the grooves and edges of the β-sheet of the fibrils, and the modeling of AlphaFold-Multimer was used for refining the results; secondly, Sequence-based design, which involved a peptide language model (PepMLM) that generated the variants coming from the conserved amyloidogenic segments[26,44].

After screening numerous generated candidates, the sequences that were T1–T4 and S1–S4, eight of the highest-ranking ones, were synthesized and put to the test. The synergistic use of Thioflavin-T (ThT) fluorescence assays and transmission electron microscopy (TEM) unveiling of not only a few peptides but also the two mentioned, T1 and S1, that had very strong dimerizing effects signaling the decline of α-syn aggregation and modifying the morphology of fibrils. These peptides were getting rid of fibrils by promoting the formation of amorphous or fragmented aggregates rather than mature amyloids.

In summary, this paper presents a methodology which can be applied in a similar way for ai-assisted peptide design. The combination of cryo-EM structural knowledge with that of deep learning–based sequence generation not only enables the easy rational thing to do for discovering the inhibitors of the conserved aggregation motifs but also the process becomes a two-way street. This technique, in conjunction with α-synuclein, can easily disseminate across the board of all amyloidogenic proteins such as tau, Aβ, and TDP-43, thus signifying a bright future in the field of drug development for the treatment of neurodegenerative diseases, via precision therapeutics.

## 2. Materials and Methods

### 2.1. Expression and purification of recombinant α-synuclein

Recombinant human wild-type α-synuclein was made in E. coli BL21(DE3) with a pET28a vector. The bacteria were grown in Luria–Bertani (LB) medium at 37 °C till the OD_600_ was 0.6 (optical density at 600 nm), then, were induced for 4 h with 1 mM IPTG (isopropyl β-D-1-thiogalactopyranoside). Cells were collected by centrifugation at 4000 rpm for 10 minutes, then, were resuspended in lysis buffer (20 mM Tris-HCl, pH 7.5, 150 mM NaCl), and were disrupted by probe sonication (3 s on / 3 s off cycles at 60% amplitude for a total of 10 minutes).

The lysate was subjected to centrifugation at 18,000 × g for 30 minutes at 4 °C for clarification. The bacterial proteins were precipitated by heat denaturation at 100 °C for 10 minutes. The supernatant was subjected to filtration (0.45 µm) and subsequently loaded onto Q-Sepharose Fast Flow anion exchange column equilibrated with buffer A (20 mM Tris-HCl, pH 7.4). The proteins attached to the column were removed by a linear NaCl gradient (0–500 mM). The fractions containing α-synuclein were combined, dialyzed against 20 mM Tris-HCl (pH 7.4), and further purified through size-exclusion chromatography (Superdex 200 Increase 10/300 GL column) with 20 mM Tris and 100 mM NaCl (pH 7.4) as buffer. Protein purity (> 95%) was confirmed by SDS-PAGE and electrospray ionization mass spectrometry. The protein concentrations were measured spectrophotometrically at 280 nm with an extinction coefficient of 5960 M^−1^ cm^−1^.

### 2.2. In vitro fibrillization of α-synuclein

The frozen monomeric α-synuclein was thawed and then subjected to ultracentrifugation (100,000 × g, 30 min, 4 °C) in order to eliminate any preformed aggregates. The preparation of aggregation reactions (100 µL) suggested the use of 100 μM α-synuclein in 20 mM Tris-HCl (pH 7.4), 150 mM NaCl, and 0.01% NaN_3_, with or without the addition of peptide inhibitors at specified molar ratios (α-syn:peptide = 1:0.5, 1:1, or 1:2). In addition, the optical film was used to seal the 96-well microplate (nonbinding surface, black walls, clear bottom) thus preventing evaporation of the samples. The reactions were continuously shaken (orbital, 600 rpm) at 37 °C for a maximum of 72 h.

In the case of seeded reactions, the preformed α-syn fibrils that were produced under identical conditions (no peptides) were sonicated to obtain short seeds (30 s pulse, 10% amplitude) and then incorporated at the rate of 1% (w/w) into the monomeric reactions. The fibril formation kinetics were tracked by means of Thioflavin-T fluorescence.

### 2.3. Identification of conserved structural motifs from cryo-EM data

High-resolution cryo-EM structures of α-syn fibrils from various sources including patient-derived, seeded, and recombinant were retrieved from the Protein Data Bank (PDB). Different polymorphs representing various disease states, namely MSA, PD, and DLB, were aligned through the use of PyMOL and UCSF ChimeraX software in order to reveal the structurally conserved β-sheet cores and the interfaces between the different protofilaments (Figure 1). Four conserved β-strand–rich areas (Sites 1–4) were identified according to the following criteria: (i) being present in more than 80% of known polymorphs; (ii) being accessible on the surface at either the ends of the fibrils or the junctions between the protofilaments; and (iii) having a high density of hydrogen bonds. These motifs that were identified as conserved then became the templates for the AI-guided generation of peptides.

**Figure 1.**
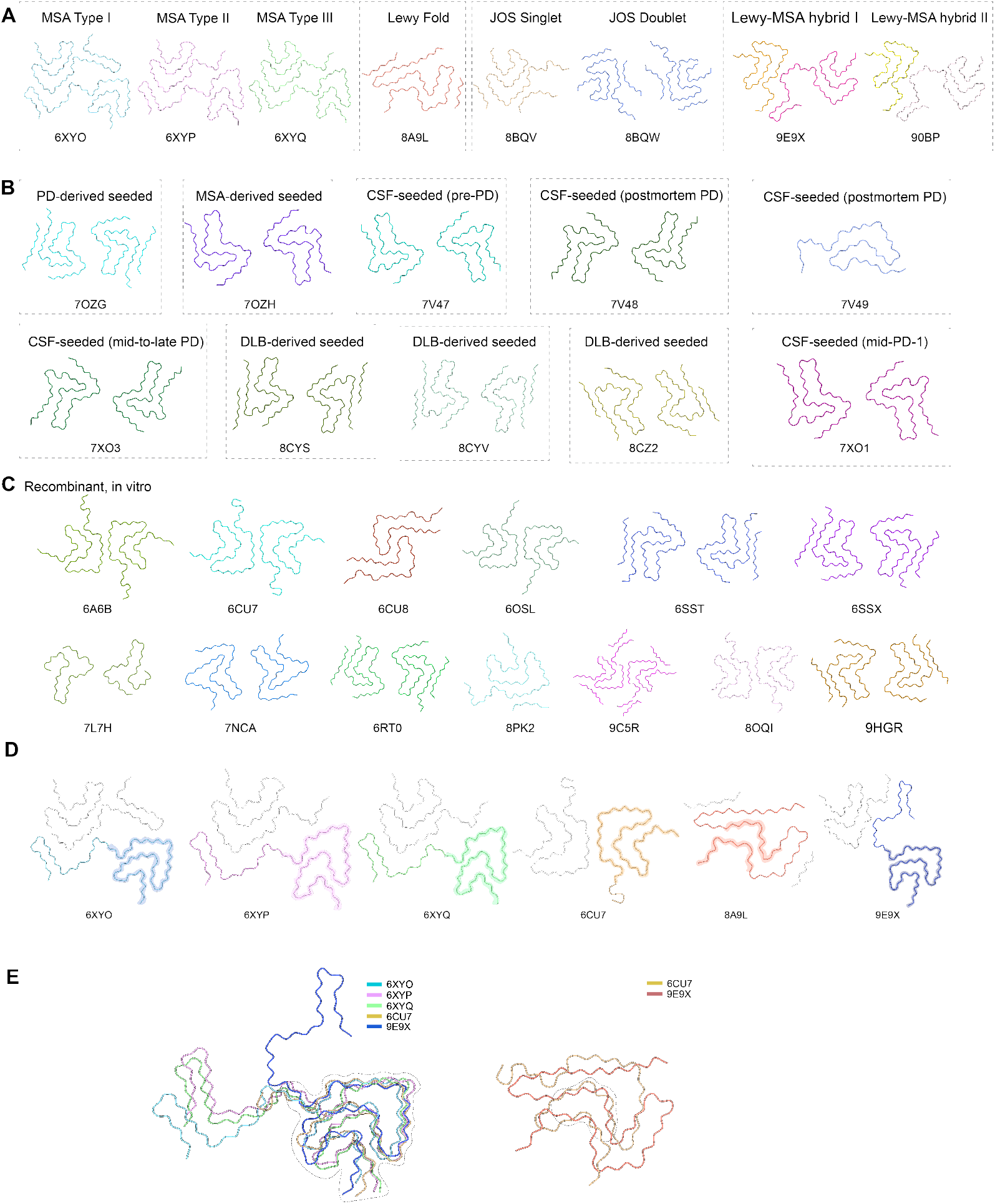
Structural diversity of α-synuclein fibrils revealed by cryo-EM. **A**. Patient-derived fibrils obtained directly from human brain tissues display distinct disease-specific conformations, including MSA type I–III (PDB: 6XYO, 6XYP, 6XYQ), Lewy body fold (PDB: 8A9L), juvenile-onset synucleinopathy singlet and doublet (PDB: 8BQV, 8BQW), and Lewy-MSA hybrid folds (PDB: 9E9X, 9QBP). **B**. Seed-amplified fibrils generated in vitro using patient-derived seeds recapitulate pathological polymorphs from Parkinson’s disease (PDB: 7OZG), MSA (PDB: 7OZH), cerebrospinal fluid–seeded samples from preclinical, postmortem, and mid-to-late-stage PD (PDB: 7V47, 7V48, 7V49, 7XO3, 7X01), as well as dementia with Lewy bodies (DLB)-derived seeds (PDB: 8CYS, 8CYV, 8CZ2). **C**. Recombinant fibrils formed from purified α-synuclein expressed in vitro reveal additional structural polymorphs, including wild-type fibrils (PDB: 6CU7, 6CU8, 6SST, 6SSX), familial mutants such as E46K (PDB: 6OSL), H50Q (PDB: 7L7H), and A53T (PDB: 7NCA), and further recombinant assemblies (PDB: 6A6B, 6RT0, 8PK2, 9C5R, 8OQI, 9HGR). **D**. Comparative structural analysis of patient-derived, seeded, and recombinant fibrils reveals both disease-specific folds and shared motifs across different conditions. **E**. Structure alignment highlights conserved β-sheet–rich core regions that are preserved across fibril types, suggesting common structural determinants underlying α-synuclein aggregation and pathology.

### 2.4. AI-guided peptide design workflow

A dual design approach consisting of both structure-based and sequence-based modeling was applied and the methodology is presented in Figure 2.

**Figure 2.**
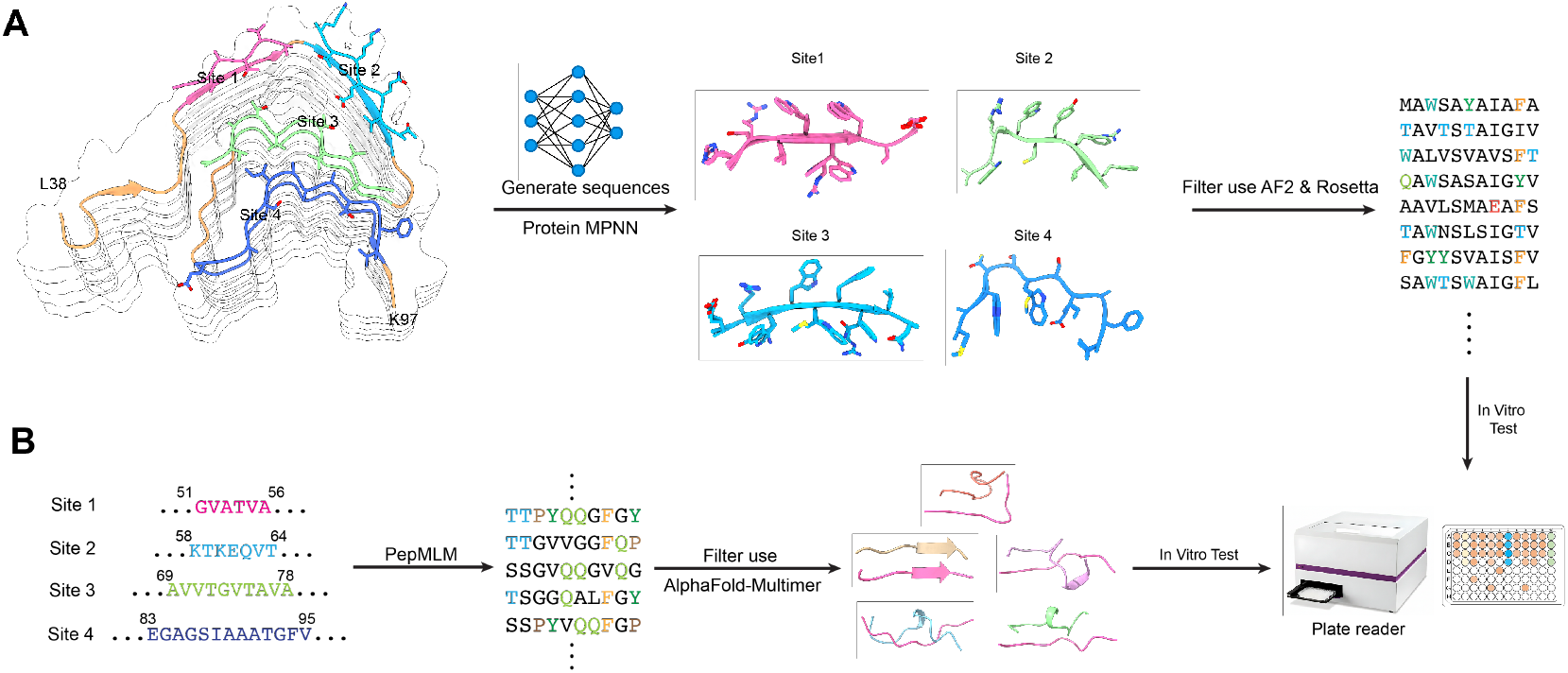
Dual design strategies for peptide inhibitors targeting conserved structural motifs of α-synuclein amyloid fibrils. **A**. Structure-based design. Conserved binding sites (Sites 1–4) on the α-synuclein fibril architecture were identified as potential capping positions to interfere with fibril elongation. ProteinMPNN was employed to generate diverse candidate peptide sequences, which were subsequently filtered and ranked using AlphaFold2 (AF2) and Rosetta to assess binding compatibility and structural stability. **B**. Sequence-based design. Conserved fibril segments were used as seed motifs to drive peptide generation via PepMLM, a sequence-based generative language model. The resulting candidates were screened using AlphaFold-Multimer to predict their binding modes with fibril segments. Shortlisted peptides from both design strategies were subjected to in vitro functional assays using a plate reader to evaluate inhibitory efficacy.

#### 2.4.1. Structure-based peptide design

For each of the conserved sites (Sites 1–4), a particular part of the fibril (6-15 residues per chain, along with neighboring β-strands) was selected. The ProteinMPNN model created numerous peptide candidates that were later assumed to help in getting the fibril interface’s backbone and side-chain geometry. Out of these, the candidates were filtered using the following criteria:

Predicted binding confidence: evaluated through AlphaFold-Multimer models which were used to predict the pLDDT and RMSD values at the interface; Binding stability: through Rosetta InterfaceAnalyzer for ΔG of binding and surface area buried; Physicochemical properties: solubility (GRAVY index < 0.5), net charge (−2≤Z≤+2), and very low aggregation propensity (TANGO < 5%). The highest-ranked candidates got the names T1–T4. Peptides with a large self-aggregate or very hydrophobic patches were not taken into account.

#### 2.4.2. Sequence-based peptide design

At the same time, the PepMLM model, which is a natural language processing model trained on natural and synthetic peptide datasets, was used to develop peptide variants from conserved amyloidogenic regions (e.g., residues 61–95 within the NAC domain). The model performed by placing hydrophilic substitutions on the solvent-exposed sites that required hydrogen-bond compatibility for solubility reasons. The sequences that were generated were admitted to the screening of: AlphaFold-Multimer for binding against fibril templates; Rosetta for scoring interface ΔG and complementarity of packing; Toxicity and aggregation predictions to eliminate potentially self-associating peptides. The top four candidates were named S1–S4 as they were designated. All lead peptides underwent further optimization for stability and solubility. Peptides with unstable secondary structures or overlap with known α-syn aggregation epitopes were excluded from further study.

### 2.5. Peptide synthesis and characterization

All candidate peptides were synthesized by solid-phase Fmoc chemistry (GenScript, USA) and purified to ≥ 95% purity by reverse-phase HPLC (C18 column). Molecular weights were confirmed by MALDI-TOF mass spectrometry. Lyophilized peptides were stored at −20 °C until use. Before experiments, peptides were dissolved in ultrapure water or minimal DMSO (< 0.1%) and diluted in buffer to final concentrations ranging from 0–140 µM. Concentrations were confirmed by amino acid analysis.

### 2.6. Thioflavin-T fluorescence assay

Amyloid formation was monitored by Thioflavin-T (ThT) fluorescence. Reaction mixtures (100 µL) contained 70 µM α-synuclein, 25 µM ThT, and peptide inhibitors at specified concentrations. Fluorescence readings were collected every 10 min using a plate reader (excitation = 440 nm; emission = 485 nm) with intermittent shaking. Increasing ThT fluorescence indicated β-sheet formation. Each experiment was performed with three independent biological replicates, each measured in triplicate. Data were presented as mean ± SD and analyzed by one-way ANOVA followed by Tukey’s multiple-comparison test.

### 2.7. Transmission electron microscopy (TEM)

To examine fibril morphology, samples collected at the end of aggregation reactions were adsorbed (10 µL) onto carbon-coated copper grids (400-mesh) for 2 min, blotted, rinsed twice with distilled water, and negatively stained with 2% (w/v) uranyl acetate for 1 min. After air drying, grids were imaged using a JEOL JEM-1400 TEM operating at 120 kV.

Micrographs were captured at 50,000× magnification and analyzed with ImageJ. For each condition, at least ten images from independent grids were quantified. Fibrils were classified as long (> 500 nm), short (100–500 nm), or amorphous (< 100 nm). The percentage of each morphology type was determined from segmented images.

### 2.8. Statistical and computational analyses

All numerical analyses were performed in Python (NumPy, Pandas, SciPy) and GraphPad Prism 9.0. Data were reported as mean ± SD. Statistical comparisons included: Aggregation kinetics: analyzed by one-way ANOVA for lag-phase and plateau differences; Morphological distributions: evaluated using chi-square tests; Computational– experimental correlation: assessed by Pearson’s correlation between predicted ΔG values and observed inhibition percentages. All computational workflows (ProteinMPNN, PepMLM, AlphaFold-Multimer, Rosetta InterfaceAnalyzer) were executed using standardized parameters to ensure reproducibility.

### 2.9. Data availability

All raw and processed ThT fluorescence data, peptide sequences, and computational scripts are available upon reasonable request to the corresponding author. Cryo-EM structural models used in this study were obtained from the Protein Data Bank under accession codes listed in Figure 1.

## 3. Results

### 3.1. Structural diversity of α-syn fibrils and identification of conserved β-sheet motifs

For the purpose of discovering universal inhibitory targets, we investigated cryo-EM structures of α-syn fibrils coming from different sources in a systematic manner, such as patient brain tissues, seeded in vitro fibrils and recombinant assemblies (Figure 1). Notwithstanding the pronounced differences in the protofilament interfaces and cross-β geometries among the various fibril types connected with multiple system atrophy (MSA), Parkinson’s disease (PD) and juvenile-onset synucleinopathy, all types still could not avoid the common hydrophobic core of residues 37–98 that correspond to the non-amyloid-β component (NAC) region.

The structural alignment of 20 representative fibril polymorphs exposed the presence of four β-strand clusters (Sites 1–4) characterized by recurrent high hydrogen-bond density and very small atomic deviation (RMSD < 1.2 Å), which is a strong indicator of structural conservation of the fibrils. While Sites 1 and 2 were mainly at the fibril ends, giving them the potential for inhibition through end-capping, Sites 3 and 4 were along the inter-protofilament interfaces, creating “cross-hair” or “Greek key” structural motifs.

The analysis of sequence conservation also pointed at recurring hydrophobic and aromatic residues (V66, V70, A76, F94) which made β-sheet contacts that are crucial for the stability of the fibril. Therefore, these four regions were designated as the design templates for the peptide inhibitor development targeting the conserved β-sheet grooves and interfacial pockets responsible for fibril growth and stabilization.

### 3.2. AI-guided dual design and in silico screening of peptide candidates

The use of the dual AI-guided pipeline resulted in the generation and screening of over 10,000 peptide candidates (6– 15 amino acids long) through a combination of structure-based (ProteinMPNN) and sequence-based (PepMLM) methods. After computational filtering of the candidates based on solubility, interface complementarity, and predicted stability, the eight top-ranked candidates T1–T4 (structure-based) and S1–S4 (sequence-based) were selected for experimental validation (Table 1 and Table 2). Predicted binding models indicated different inhibitory mechanisms for the different candidates (Figure 2). T1 and T2 peptides had their binding longitudinally along β-sheet edges, forming parallel backbone hydrogen bonds that could sterically hinder the addition of the monomer. T3 and T4 interacted perpendicularly across the protofilament interfaces, which may lead to the destabilization of lateral packing. S1 and S2 were mimetic of short NAC fragments, but they had hydrophilic substitutions at solvent-exposed positions, thus improving solubility while still being compatible with the β-sheet. S3 and S4 were constructed to develop amphipathic helices that would be able to cap fibril ends through hydrophobic insertion.

**Table 1.**
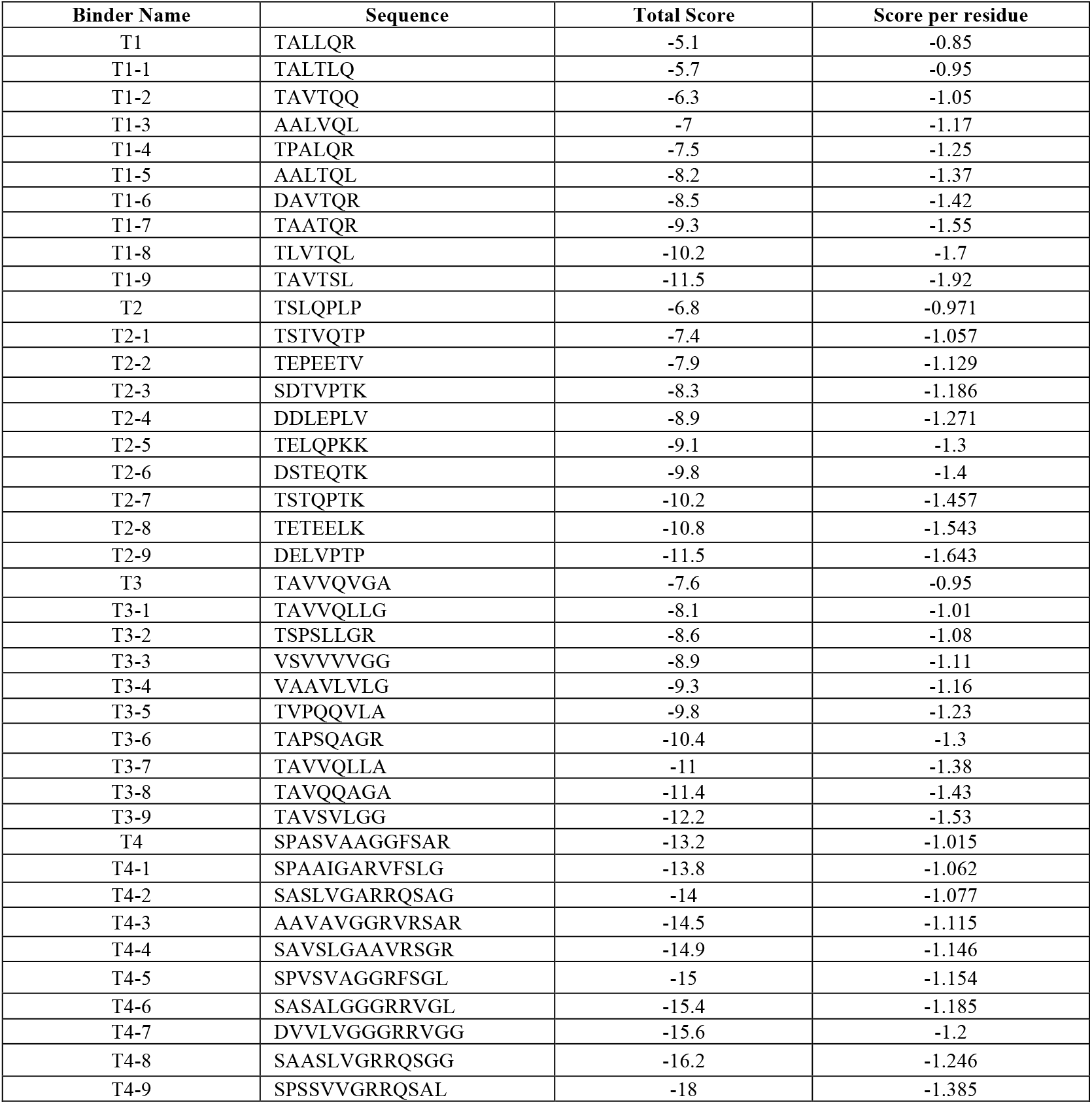
Peptide Sequences, Total Scores, and Scores Per Residue Generated by ProteinMPNN.

**Table 2.**
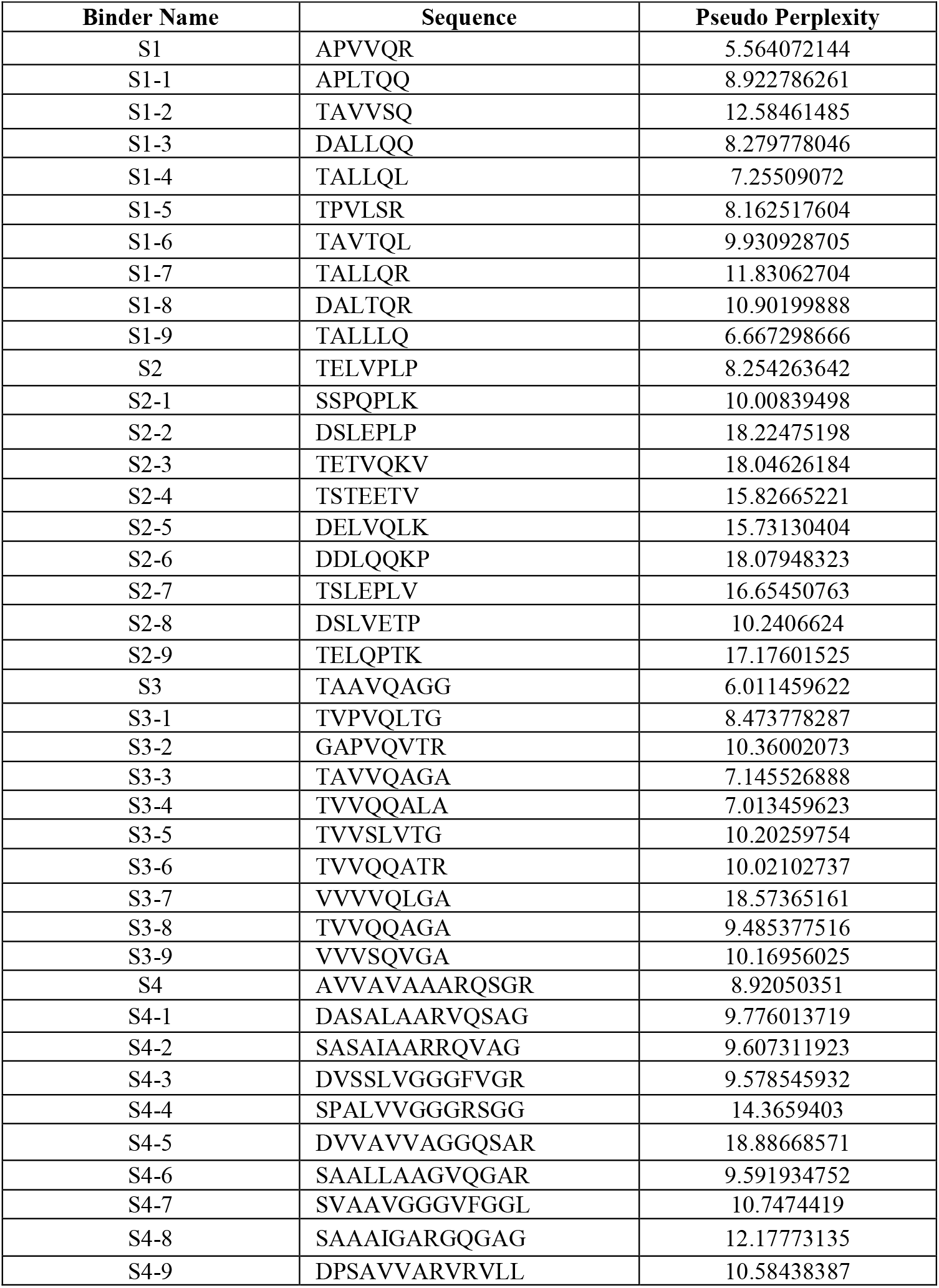
Peptide Binder Sequences and Pseudo Perplexity from PepMLM.

Energetic scoring with Rosetta InterfaceAnalyzer gave binding free energies (ΔG) of between −24 and −36 kcal/mol for the leading candidates, while AlphaFold-Multimer predicted interface confidence scores (pTM > 0.6) for T1, T2, and S1 complexes. Such results gave a strong computational backing for the assumption of their high binding affinity and structural compatibility, which in turn, led to informed prioritization of subsequent experimental work.

### 3.3. Inhibition of α-syn aggregation monitored by Thioflavin-T fluorescence

For testing the effect of inhibition, Thioflavin-T (ThT) fluorescence kinetics were taken as a measure for α-synuclein aggregation in the presence or absence of each of the designed peptide (Figure 3). In the control experiments that only consisted of α-synuclein monomers, the typical sigmoidal aggregation curves had shown up, which were marked by a lag phase (∼16 h), followed by an exponential rise, and then a fluorescence plateau after ∼60 h.

**Figure 3.**
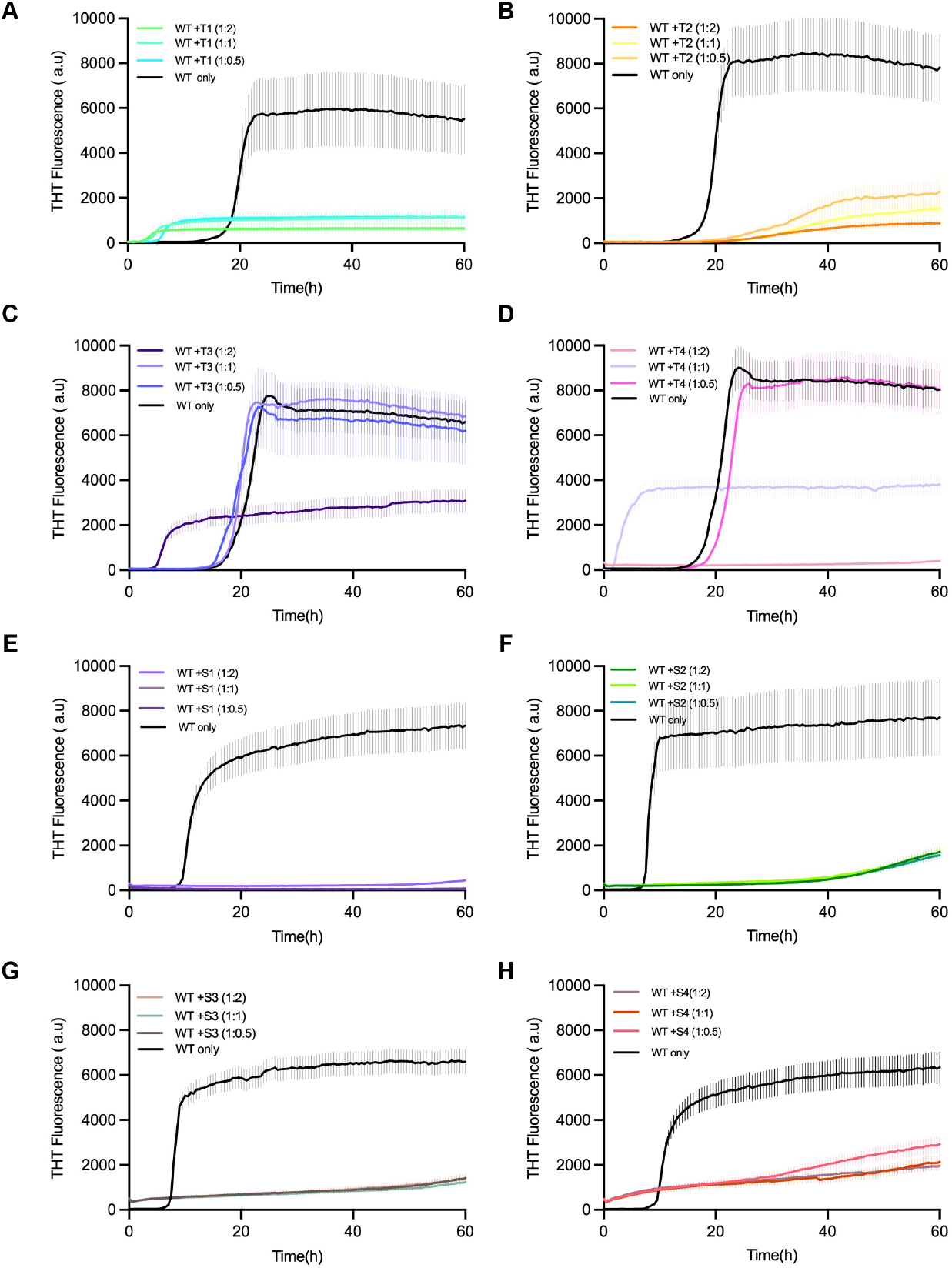
Inhibition of α-synuclein fibril formation by designed linear peptides T1–T4 and S1–S4. Thioflavin T (ThT) fluorescence assays showing the aggregation kinetics of α-synuclein monomers in the presence of designed linear peptide inhibitors. (**A–D**) Correspond to T1–T4 peptides; (**E–H**) correspond to S1–S4 peptides. α-Synuclein monomers (70 µM) were incubated alone (black) or with each peptide at molar ratios of 1:0.5, 1:1, and 1:2 (monomer:peptide). ThT fluorescence was monitored over 60 h to assess amyloid fibril formation. All peptides exhibited inhibitory effects on α-synuclein aggregation to varying degrees, reflected by reduced fluorescence intensity and prolonged lag phases compared to WT-only controls. Data are presented as mean ± s.d. from three independent replicates.

On the other hand, all the reactions treated with the peptides showed that the aggregation onset was delayed and the final ThT intensity was diminished, indicating the effective inhibition of β-sheet formation. Of the peptides that were tested, T1 and S1 had the most powerful inhibitory action, almost completely eliminating fluorescence at a 1:2 α-syn:peptide molar ratio. Peptides T2 and S2 provided moderate inhibition, while T3–T4 and S3–S4 resulted in minor suppression.

The decline in fluorescence that was dose-dependent confirmed that the inhibition was connected with the concentration of peptide. Thus, it can be interpreted that the designed peptides are able to interfere with α-syn fibrillation in a concentration-dependent manner, and this, in turn, leads to successful disruption of amyloid propagation.

### 3.4. Morphological modulation by peptide inhibitors

Negatively stained transmission electron microscopy (TEM) was used as a tool to visualize fibril morphology and it was done at the end of the aggregation reactions (Figure 4). The untreated control formed a dense network of long, twisted fibrils which averaged about ∼800 ± 150 nm in length and ∼10 ± 2 nm in diameter.

**Figure 4:**
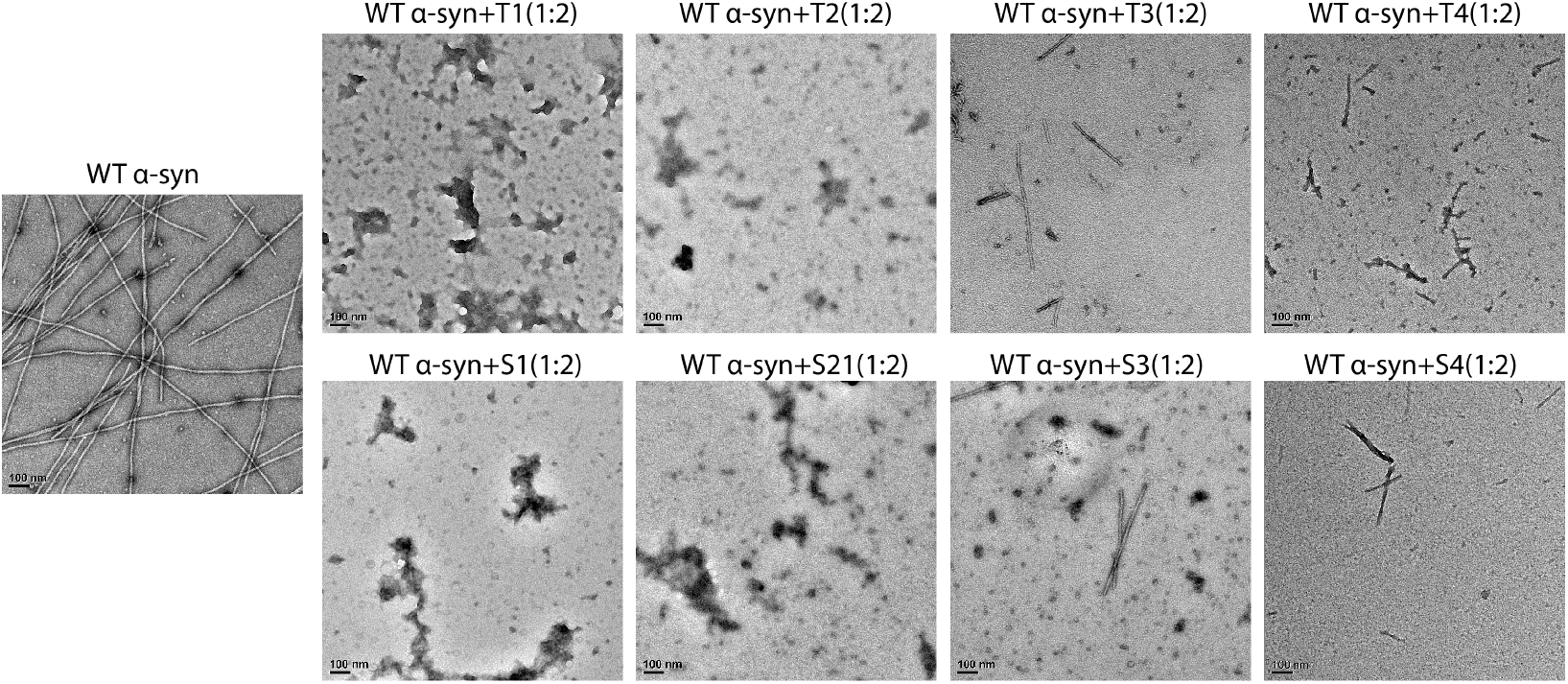
Morphological modulation of wild-type α-synuclein (α-syn) fibrillation by designed peptide inhibitors. Transmission electron microscopy (TEM) negative stain images show the aggregation morphology of wild-type α-syn (WT α-syn) with and without treatment with T-series and S-series peptide inhibitors. The WT α-syn control group shows highly mature and well-defined α-syn amyloid fibrils, with typical morphological features. All inhibitor-treated groups were at a molar ratio of 1:2 (inhibitor: α-syn monomer). T1 and S1 inhibitors showed the strongest anti-aggregation activity at the tested concentration, leading to the complete disappearance of mature fibrils, with products mainly consisting of amorphous or small granular aggregates. This suggests they effectively block fibril elongation and stabilization. T2 and S2 also significantly reduced the number and length of fibrils, but a few scattered rod-like or fragmented aggregates could still be observed. In contrast, the inhibitory effects of T3, T4, S3, and S4 were relatively weaker, with a number of short fibrils and incompletely disassembled intermediates still visible. This indicates that the T1 and S1 peptides have a high degree of morphological inhibitory effect on α-syn aggregation. The scale bar in all images is 100 nm.

On the other hand, the samples that were treated with T1 or S1 had almost no mature fibrils present, just some amorphous or short fragmented aggregates (< 100 nm) left behind. T2 and S2 treatments caused a significant reduction in the number of fibrils, with mainly short fragments (200–400 nm) and a lower overall density of fibrils. The treatments of T3–T4 and S3–S4 resulted in the formation of mixed populations of short fibrils and disordered aggregates.

The quantitative image analysis verified that there was a reduction of approximately 90% of the long fibrils in the T1/S1 groups when compared to the control (p < 0.001). Additionally, the fibril density had a strong positive correlation with the ThT fluorescence intensity (Pearson’s r = 0.91), which indicates the consistency between the fluorescence and TEM results. The combined findings suggested that the peptides, and especially T1 and S1, not only hindered fibril nucleation but also diverted aggregation pathways towards the non-fibrillar assemblies.

### 3.5. Correlation between computational predictions and experimental inhibition

The predictive accuracy of the AI-guided design was evaluated by comparing computational parameters—Rosetta binding free energy (ΔG) and AlphaFold interface confidence (pTM)—with experimental inhibition efficiencies. A high degree of correlation was found (r = 0.87, p < 0.01), and T1 and S1 were the compounds with the most favorable predicted binding metrics as well as the highest experimental inhibition, indicating a good correlation between the predictions and the experimental results.

The AI-assisted design pipeline was confirmed as predictive and reliable, indicating that structural modeling combined with machine learning–driven sequence generation can successfully prioritize biologically active inhibitors before their synthesis.

## 4. Discussion

The present study sets up a framework for the rational design of peptide inhibitors against α-synuclein (α-syn) fibril polymorphs that is based on structure and informed methods. We discovered small peptides that greatly inhibit α-syn aggregation in vitro by integrating cryo-electron microscopy (cryo-EM)–derived structure data with computational sequence generation. Notably, although the different conformational shapes of α-syn fibril polymorphs are very different from each other, they all have one common feature, i.e., the β-sheet motifs that are conserved and stabilized by interactions of hydrophobic and backbone hydrogen-bond types. Such features that occur frequently allow for the creation of constitutively conserved binding sites across different fibril strains that can be utilized for broad inhibition. The experimental results suggest that the peptides designed have two major pathways through which they affect α-syn aggregation. For instance, T1 and S1 peptides are seen to attach themselves to the ends of the fibrils, thus, capping the β-sheet edges and blocking the addition of new monomers. Others like T2 and S2 are presumed to bind at the inter-protofilament contact points; thereby, diminishing the contacts and facilitating the separation of the f.t.n. The reduction in Thioflavin-T-fluorescence and the vanishing of the older fibrils in the transmission electron microscopy images support these mechanisms. The different but concurrent mechanisms imply that the use of conserved β-sheet regions as a target is a reliable strategy to successfully cut off the propagation of the fibrils regardless of their polymorphic variation.

One of the major advantages of this method was the complementary approach of structure-based and sequence-based design pipelines. The structure-based technique, which was supported by ProteinMPNN and AlphaFold-Multimer, guaranteed geometric and energetic complementarity with atomically detailed templates, while the sequence-based model (PepMLM) allowed for the wider exploration of sequence space beyond the known amyloidogenic motifs. The most potent inhibitors, T1 and S1, were, importantly, obtained through different but coalescent computational techniques, thus emphasizing the blending of orthogonal design paradigms. The excellent agreement between computational predictions and experimental potency is an additional proof that binding energy and confidence metrics can be used as trustworthy pre-screening criteria, thus cutting down on the experimental workload and enhancing design efficiency.[45,46].

From the viewpoint of therapy, the aggregation of α-syn is the major process in the development of Parkinson’s disease (PD) and essentially all types of synucleinopathies[20,21,47]. During the current approaches using small molecules and antibodies, the attempts to therapeutic intervention have not been very effective, in part, due to the many different conformations of the α-syn aggregates[30]. The present work on the peptide inhibitors has the possibility to target the structural features which are universally present in all polymorphs thus being a potential solution for overcoming the issue[21]. The peptide’s unique characteristics of being modular and having adjustable chemistry make them eligible for additional improvements in terms of stability, brain permeability, and resistance to metabolizing by using backbone cyclization, D-amino acid substitution, or nanoparticle conjugation.

This study has several limitations that must be acknowledged. All tests were performed in vitro, which are conditions that do not entirely mimic the intracellular environment. The work planned for the future should check if the peptides work against α-syn aggregation and toxicity in neuronal and in vivo models. It will be necessary to carry out high-resolution structural characterization of peptide fibril complexes using methods like cryo-EM or solid-state NMR to get the binding modes confirmed and the mechanistic understanding refined. Moreover, kinetic analyses separating the nucleation and elongation rate constants could assist in identifying the stage-specific inhibitory effects. Lastly, pharmacokinetic profiling and AI-assisted stability and delivery optimization will be indispensable to getting these peptides ready for therapeutic use.

## 5. Conclusions

This study discloses a combined computational-experimental method for designing peptide inhibitors which will, more or less, target α-syn fibril polymorphs. The integration of cryo-EM structural insights with algorithm-guided sequence design enabled the detection of short peptides that could severely moderate α-syn fibrillation and change fibril morphology in vitro. The best-performing ones, T1 and S1, almost wholly prevented amyloid formation through terminal capping and interfacial disruption mechanisms.

The given evidence points out that the fusion of structural biology with computational peptide design quickens and makes rational the discovery of aggregation inhibitors. The methodology developed at this point is applicable to other amyloidogenic proteins like tau, Aβ, and TDP-43 and lays the groundwork for precision-engineered therapeutics that counteract protein misfolding in neurodegenerative diseases.

## Author Contributions

J.D., H.Z., and C.S. designed and performed the research. J.D., H.Z., and C.S. contributed to methodology development, data acquisition, and analysis. Conceptualization and manuscript preparation were jointly carried out by J.D., H.Z., and C.S. under the supervision of C.S. Data curation was performed by C.S. Funding was secured by L.J. All authors reviewed and approved the final version of the manuscript.

## Institutional Review Board Statement

Not applicable. No human or animal subjects were involved in this study.

## Informed Consent Statement

Not applicable.

## Data Availability Statement

All data supporting the findings of this study are available from the corresponding author upon reasonable request. Cryo-EM models analyzed in this work were obtained from the Protein Data Bank, with accession codes listed in Figure 1.

## Acknowledgments

The authors thank Dr. Lin Jiang for generous financial support. The electron imaging was performed using the facilities of the Electron Imaging Center for Nanomachines (EICN) at the California NanoSystems Institute (CNSI), UCLA. The authors also thank members of the Jiang Laboratory for valuable discussions and technical assistance.

